# Shared neural underpinnings of multisensory integration and trial-by-trial perceptual recalibration

**DOI:** 10.1101/566927

**Authors:** Hame Park, Christoph Kayser

## Abstract

Multisensory stimuli create behavioral flexibility, e.g. by allowing us to derive a weighted combination of the information received by different senses. They also allow perception to adapt to discrepancies in the sensory world, e.g. by biasing the judgement of unisensory cues based on preceding multisensory evidence. While both facets of multisensory perception are central for behavior, it remains unknown whether they arise from a common neural substrate. In fact, very little is known about the neural mechanisms underlying multisensory perceptual recalibration. To reveal these, we measured whole-brain activity using MEG while human participants performed an audio-visual ventriloquist paradigm designed to reveal multisensory integration within a trial, and the (trial-by-trial) recalibration of subsequent unisensory judgements. Using single trial classification and behavioral modelling, we localized the encoding of sensory information within and between trials, and determined the behavioral relevance of candidate neural representations. While we found neural signatures of perceptual integration within temporal and parietal regions, of these, only medial superior parietal activity retained multisensory information between trials and combined this with current evidence to mediate perceptual recalibration. These results suggest a common neural substrate of sensory integration and trial-by-trial perceptual recalibration, and expose the medial superior parietal cortex as a flexible hub that links present and previous evidence within and between senses to guide behavior.

## Introduction

Multisensory information offers substantial benefits for behavior. For example, acoustic and visual cues can be combined to derive a more reliable estimate of where an object is located (1–4). Yet, the process of multisensory perception does not end once an object is removed. In fact, multisensory information can be exploited to calibrate subsequent perception in the absence of external feedback (5,6). In a ventriloquist paradigm, for example, the sight of the puppet and the actor’s voice are combined when localizing the speech source, and both cues influence the localization of subsequent unisensory acoustic cues, if probed experimentally (7–13). This trial-by-trial recalibration of perception by previous multisensory information has been demonstrated for spatial cues, temporal cues, and speech signals (14–17). Despite the importance of both facets of multisensory perception for adaptive behavior – the combination of information within a trial and the trial by trial adjustment of perception – it remains unclear whether they originate from shared neural mechanisms.

In fact, the neural underpinnings of trial-by-trial recalibration remain largely unrevealed. Those studies that have investigated neural correlates of multisensory recalibration mostly focused on the adaptation following long-term (that is, often minutes of) exposure to consistent multisensory discrepancies (18,19). However, we interact with our environment using sequences of actions dealing with different stimuli, and thus systematic sensory discrepancies as required for long-term effects are possibly seldom encountered. Hence, while the behavioral patterns of multisensory trial-by-trial recalibration are frequently studied (6,7,9,20,21) it remains unclear when and where during sensory processing their neural underpinnings emerge.

In contrast to this, the neural underpinnings of multisensory integration of simultaneously received information have been investigated in many paradigms and model systems (22–24). Studies on spatial ventriloquist-like paradigms, for example, demonstrate contributions from auditory and parietal cortex (9,11,25–29). A series of recent studies demonstrates that posterior parietal regions automatically fuse multisensory information, while anterior parietal regions give way to a more flexible spatial representation that follows predictions from Bayesian causal inference (30–34). Given that parietal regions also contribute to the maintenance of sensory information within or between trials (35–40) this raises the possibility that parietal regions are in fact mediating both the combination of sensory information within a trial, and the influence of such an integrated representation on guiding subsequent adaptive behavior.

To link the neural mechanisms underlying multisensory integration and trial-by-trial recalibration, we measured whole-brain activity using magnetoencephalography (MEG) while human participants performed a spatial localization task. The paradigm was designed to reveal the behavioral correlates of audio-visual integration (i.e. the ventriloquist effect, VE) and the influence of this on the localization of a subsequent unisensory sounds (the ventriloquist aftereffect, VAE) (6). Using single-trial classification we determined the relevant neural representations of auditory and visual spatial information and quantified when and where these are influenced by previous sensory evidence. We then modelled the influence of these candidate neural representations on the participant-specific trial-by-trial response biases. As expected based on previous work, our results reveal neural origins of sensory integration in superior temporal and parietal regions. Importantly, of these, only activity within the superior parietal cortex encodes current multisensory information and retains information from preceding trials, and uses both to guide adaptive behavior within and across trials.

## Results

### Behavioral Results – Ventriloquist effect

Behavioral responses in audio-visual (AV) trials revealed a clear ventriloquist bias, whereby the visual stimulus biased the perceived sound location (Figure 1B). Model comparison revealed that both stimuli had a significant influence on the participants’ responses (relative BIC values of the three models: mi_1_: 938, mi2: 3816, mi3: 0; relative AIC values; mi_1_: 945, mi_2_: 3823, mi_3_: 0, winning model: mi_3_: VE ~ 1 + β·A_n-1_ + β·V_n-1_), with significant contributions from both the acoustic (A_n-1_), and visual stimuli (V_n-1_) (β_An-1_ = −0.48, β_Vn-1_ = 0.22, t_An-1_ = −70.0, t_Vn-1_ = 31.7, p_An-1_, p_Vn-1_ < 0.01, d.f. = 8064). Across participants, the VE bias was significant for each non-zero audio-visual discrepancy (all p < 10^−4^; Wilcoxon signed rank tests, corrected for multiple tests with the Holm procedure).

**Figure 1.**
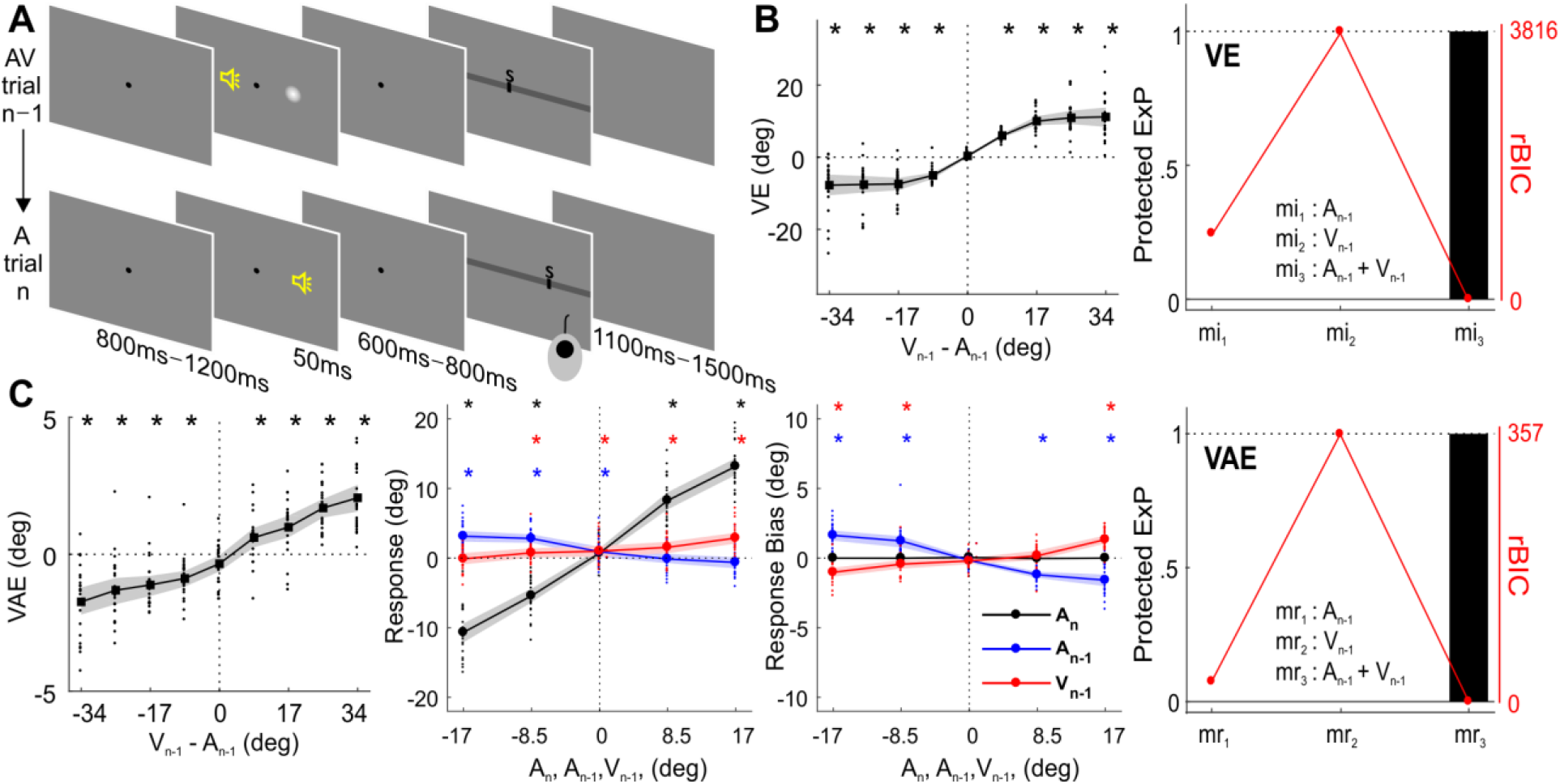
Paradigm and behavioral results (N = 24). **(A)** Experimental design. Participants localized auditory (or visual) targets and indicated the perceived location using a mouse cursor. Audio-visual (AV) and auditory (A) trials alternated. **(B) (left)** Response bias induced by the ventriloquist effect (VE) as a function of audio-visual discrepancy in trial n-1. **(right)** Relative BIC values and protected exceedance probabilities obtained by comparing different candidate models describing the VE. **(C) (left)** Response bias induced by the ventriloquist aftereffect in trial n (VAE). **(middle two)** Sound localization response and bias in trial n were significantly influenced by the current sound (A_n_), the previous sound (A_n−1_) and the previous visual (V_n-1_) stimulus. **(right)** Comparison of candidate models for the VAE. **(B-C)** Solid lines indicate mean across participants. Shaded area is the estimated 95% confidence interval based on the bootstrap hybrid method. Dots denote individual participants. ExP: exceedance probability. rBIC: relative BIC. An: sound location in trial n. A_n-1_: sound location in trial n-1. V_n-1_: visual location in trial n-1. Asterisks denote p-values < 0.05 from two-sided Wilcoxon signed rank tests corrected with the Holm procedure.

### Behavioral Results – Ventriloquist aftereffect

Behavioral responses in auditory trials revealed a significant ventriloquist aftereffect (Figure 1C), confirming that the previous multisensory stimuli had a lasting influence on the localization of subsequent sounds. Model comparison revealed that both previous stimuli had a significant influence on the VAE (relative BIC values of the three models; mr_1_: 27, mr_2_: 357, mr_3_: 0; relative AIC values: mr_1_: 34, mr_2_: 364, mr_3_: 0, winning model: mr_3_: VAE ~ 1 + β·A_n-1_ + β·V_n-1_), with significant contributions from the previous sound (A_n-1_), and the previous visual stimulus (V_n-1_) (β_An-1_ = −0.09, β_Vn-1_ = 0.03, t_An-1_ = −19.4, t_Vn-1_ = 6.0, p_An-1_, p_Vn-1_ < 0.01, d.f. = 8064). Across participants, the VAE bias was significant for each non-zero audio-visual discrepancy (all p < 10^−2^; two-sided Wilcoxon signed rank tests, corrected for multiple tests with the Holm procedure).

We performed two control analyses to further elucidate the basis of the VAE. First, we asked whether the shift in the perceived sound location was the result of a bias towards the previous visual stimulus location, or a bias induced specifically by the previous audio-visual discrepancy (6). To dissect these hypotheses, we selected trials for which the expected biases arise from the direction of the VE, and not because of a visual bias towards V_n-1_ (Figure 2A). The data were clearly in favor of a genuine multisensory bias, as the VAE remained the same and significant for these trials (all p < 0.05; except for +25.5 condition; two-sided Wilcoxon signed rank tests, corrected for multiple tests; Figure 2B)(6). Second, we asked whether the response bias in trial n was better accounted for by the sensory information in the previous trial (i.e. the previous multisensory discrepancy: Δ_n-1_ = V_n-1_ – A_n-1_) or the participant’s response in that trial (R_n-1_). Formal model comparison revealed that the model R_n_ ~ 1 + A_n_ + Δ_n-1_ provided a better account of the data than a response-based model R_n_ ~ 1 + A_n_ + R_n-1_ (relative BIC: 0, 393; BIC weights: 1, 0), supporting the notion that recalibration is linked more to the physical stimuli than the participants response (21).

**Figure 2.**
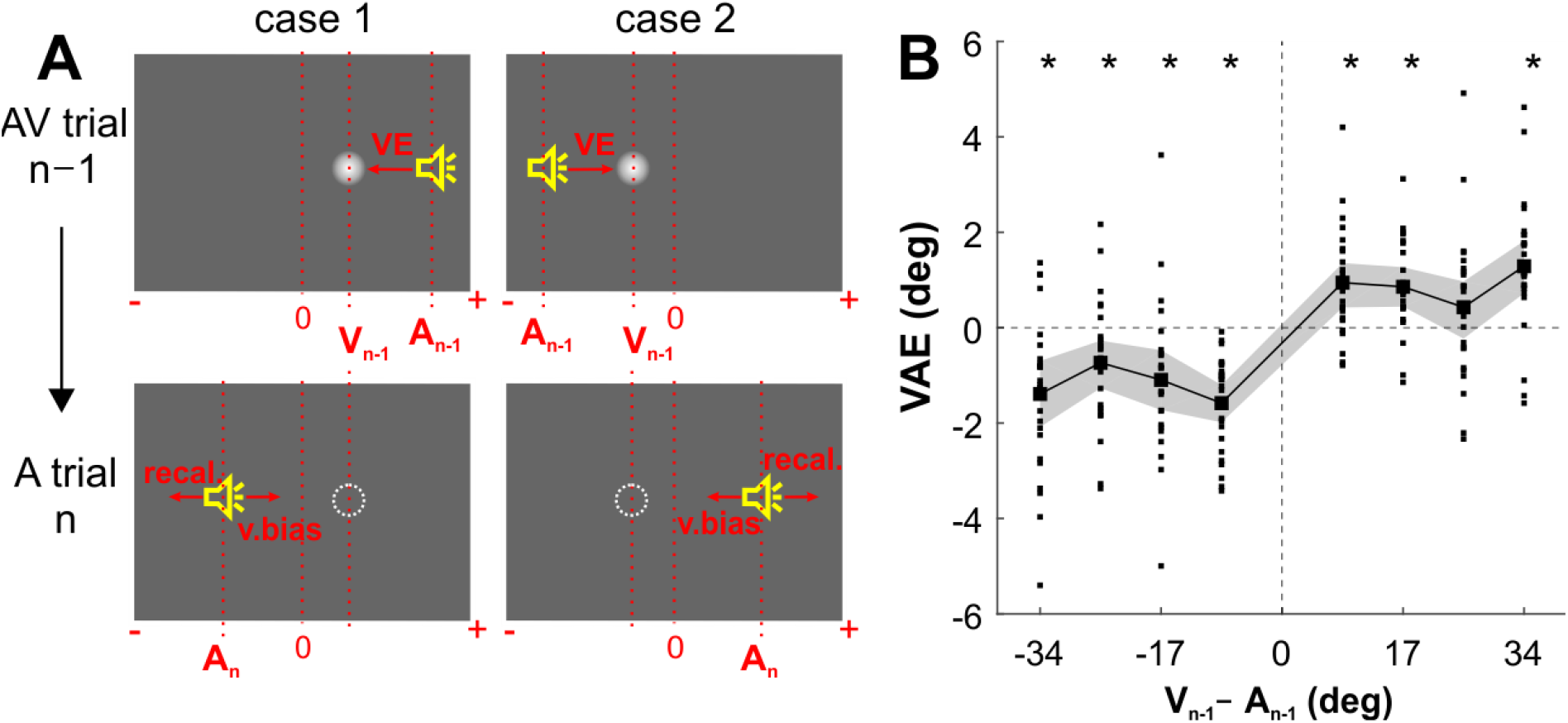
Dissociating a pure visual bias from a genuine multisensory bias in the VAE. **(A)** For this analysis we selected trials for which the expected visual (v.bias) and multisensory (recal) biases are in opposite directions. In particular, for the selected trials we would expect the VAE bias to point in the same direction as the VE bias in trial n-1, while the visual bias would be in the opposite direction (and proportional to V_n-1_ – A_n-1_). Specifically, we selected trials satisfying either case 1: V_n-1_ – A_n-1_ < 0 & A_n_ ≤ V_n-1_ or case 2; V_n-1_ – A_n-1_ > 0 & A_n_ ≥ V_n-1_. **(B)** VAE bias for these trials. Solid line indicates mean across participants. Shaded area is the estimated 95% confidence interval based on the bootstrap hybrid method. Dots denote individual participants. Asterisks denote p-values < 0.05 from two-sided Wilcoxon signed rank tests, corrected with the Holm method.

### Representations of single trial sensory information in MEG source data

The analysis of the MEG data was designed to elucidate the neural underpinnings of the VAE and to contrast these to the neural correlates of the VE. Specifically, we first determined neural representations of the task-relevant sensory information, or of the upcoming participant’s response. We then used these representations in a neuro-behavioral analysis to probe which neural representations of acoustic or visual spatial information are directly predictive of the participant-specific VE and VAE single trial biases.

We applied linear discriminant analysis to the time-resolved MEG source data to determine neural representations of the lateralization of the auditory and visual stimuli (Figure 3). From the MEG activity during the A trials we obtained significant classification performance for the current sound (A_n_; peaking at 80 ms in the left inferior parietal and at 160 ms in the middle temporal gyrus) and the location of the sound in the previous trial (A_n-1_; peaking around 120 ms in the left middle occipital lobe and the bilateral precuneus; at p ≤ 0.01 FWE corrected for multiple comparisons in source space). This characterizes neural representations of acoustic spatial information currently received and persisting from the previous trial in a wider network of temporal and parietal brain regions. Classification of the lateralization of previous visual stimuli (V_n-1_) was not significant at the whole brain level, suggesting that persistent visual information was weaker than that of the respective acoustic information. However, the whole brain classification maps revealed meaningful clusters in early left inferior temporal areas and the right inferior/superior parietal areas. Classification of the upcoming response (R_n_) was significant with a similar pattern as observed for the current sound (A_n_).

**Figure 3.**
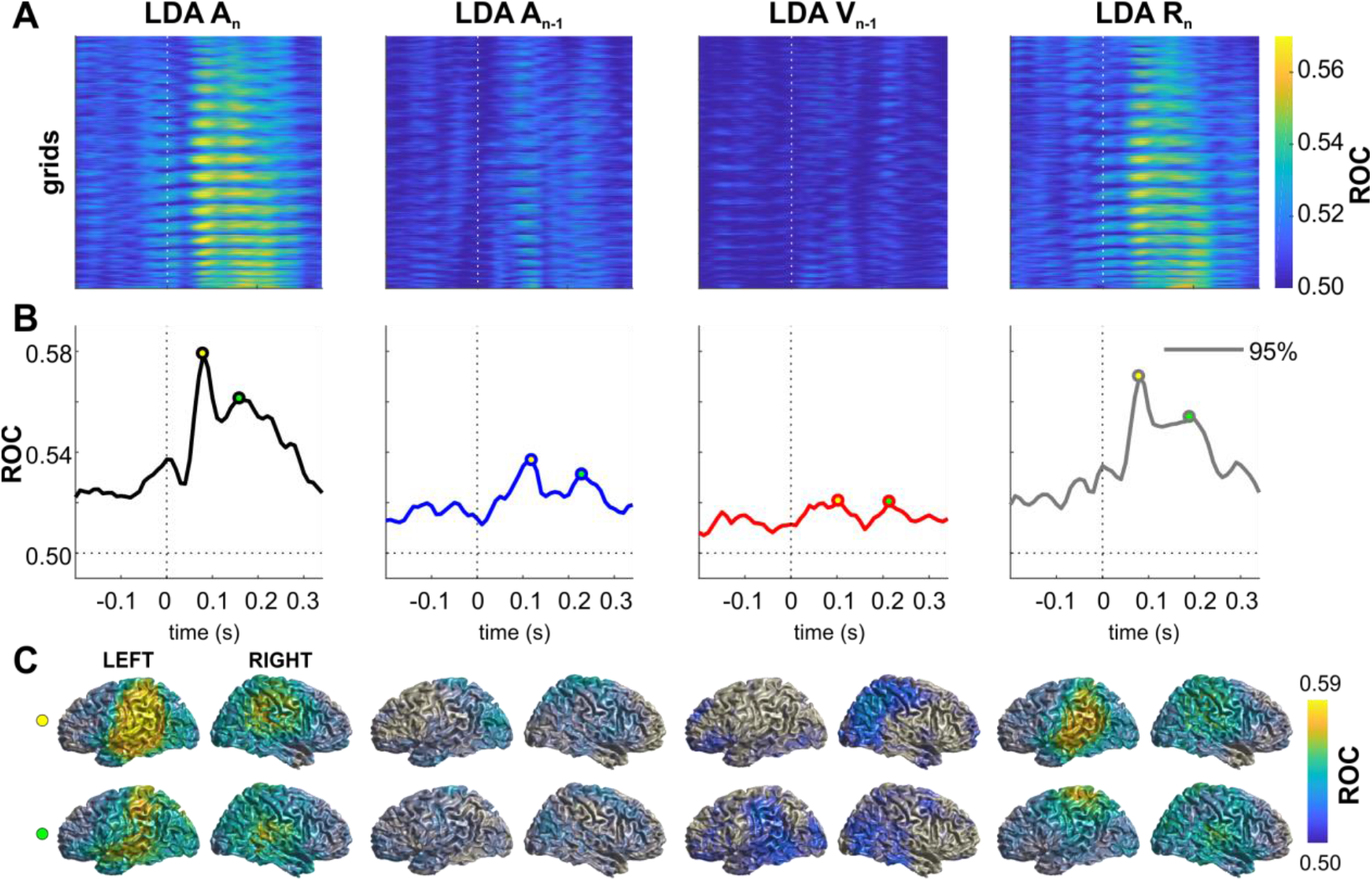
Neural representation of current and previous sensory information and upcoming responses. The figure shows the performance (ROC) of linear discriminants for different variables of interest. **(A)** Time-course of discriminant performance for all points in source space. **(B)** Time-course of the 95th percentile across source locations. **(C)** Surface projections of significant (p ≤ 0.01; FWE corrected across multiple tests using cluster-based permutation) performance at the peak times extracted from panel B (open circles). The performance of LDA V_n-1_ was not significant when tested across all source locations, and the maps for V_n-1_ are not masked with significance.

### Neural correlates of the VAE

To reveal the neural correlates of the VAE we investigated three regression models capturing different aspects of how current and previous sensory information shape i) the neural encoding of current sensory information (i.e. A_n_), ii) the encoding of the upcoming response (R_n_), and iii) the crossmodal single trial VAE recalibration bias induced.

The first model tested how the previous stimuli affect the encoding of the current sound (Figure 4A; Table 1). There was a significant (p ≤0.05; FWE corrected) influence of A*n-1*, starting around 80 ms in the cingulum, precuneus, shifting towards inferior/superior parietal areas around 220 ms. There was also significant influence of V_n-1_ in the left occipital/parietal areas around 160 ms. Importantly, the significant effects from the previous acoustic and visual stimuli overlapped in the left parietal areas (Figure 4A; red inset). The second model revealed that the previous stimuli also influenced neural activity discriminative of the participants’ response (R_n_; Figure 4B; Table 2). In particular, both previous stimuli influenced activity predictive of the response around 80 ms in the right parietal cortex (precuneus in particular), with the effect of A_n-1_ including also frontal and temporal regions. The significant effects of A_n-1_ and V_n-1_ overlapped in the cingulum and precuneus (Figure 4B; red inset). These results demonstrate that parietal regions hold information about previous multisensory stimuli, and this information affects the neural encoding of the currently perceived sound.

**Figure 4.**
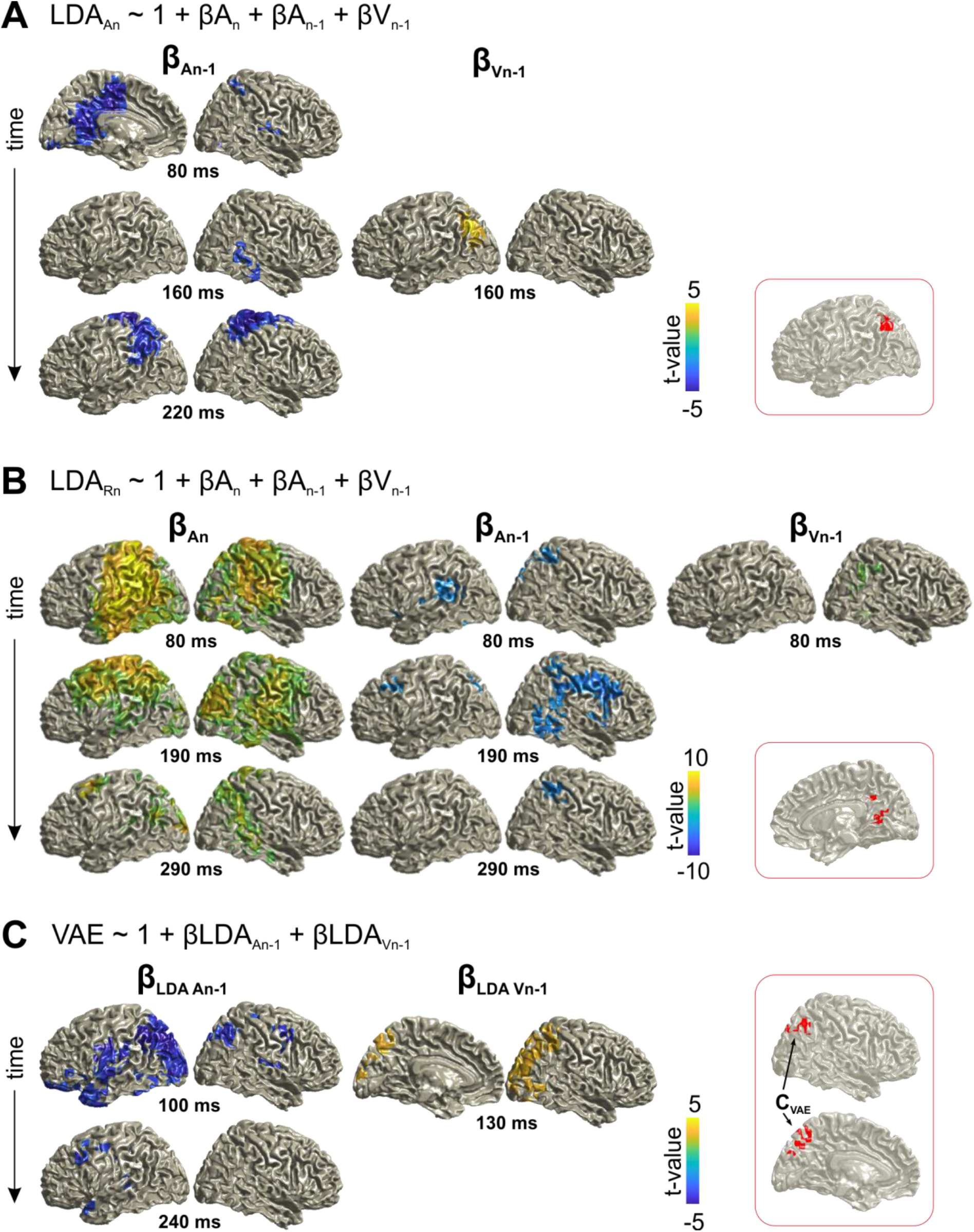
Neural correlates of trial-by-trial recalibration (VAE bias). **(A)** Contribution of previous stimuli to the neural representation of the sound (A_n_) in trial n (here the effect for A_n_ is not shown; c.f.Table 1). **(B)** Contribution of current and previous stimuli to the neural representation of the response (R_n_) in trial n (Table 2). **(C)** Ventriloquist–aftereffect bias in trial n predicted by the neural representation of information about previous stimuli (Table 3). Red insets: Grid points with overlapping significant effects for both A_n-1_ and V_n-1_ across time. Surface projections were obtained from whole-brain statistical maps (at p ≤ 0.05, FWE corrected).

**Table 1.** Regression Model: LDA_An_ ~ 1 + A_n_ + A_n-1_ + V_n-1_. The significance of each predictor was tested at selected time points at the whole-brain level (p ≤ 0.05, FWE corrected). The table provides the peak coordinates of significant clusters, the anatomical regions contributing to significant clusters (based on the AAL Atlas), as well as beta and cluster-based t-values (df = 23). The overlap was defined as grid points contributing to both a significant effect for A_n-1_ and V_n-1_ (at any time). The effect of A_n_ is not indicated, as this was significant for a large part of the temporal and parietal lobe. L: left hemisphere; R: right hemisphere. BA: Brodmann area.

| Regressor | Post-stim.<br>time (ms) | Anatomical Labels | MNI Coord. (peak) | $\beta$ t-value |
| --- | --- | --- | --- | --- |
| | | | Brodmann Area | ($t_{sum}$) |
| $A_{n-1}$ | 80 | L/R: Cingulum Mid., Precuneus | -3, -20, 29 | -5.6 |
|  |  | L: Supp. Motor Area | BA 23 | (-1220) |
|  | 160 | R: Fusiform, Temporal Mid/Inf | 28, -39, -20 | -3.8 |
|  |  |  | BA 37 | (-203) |
|  | 220 | L/R: Postcentral |  |  |
|  |  | L: Parietal Inf/Sup | -24, -36, 77 | -6.6 |
|  |  | R: Precentral, Supp. Motor Area | BA 03 | (-1560) |
| $V_{n-1}$ | 160 | L: Occipital Mid/Sup, Parietal Inf/Sup | -40, -76, 37 | 5.3 |
|  |  |  | BA 19 | (246) |
| overlap | - | L: Angular, Parietal Inf., Occipital Mid. | -40, -62, 47 | - |
|  |  |  | BA 39 |  |

**Table 2.** Regression Model: LDA_Rn_ ~ 1 + A_n_ + A_n-1_ + V_n-1_. The significance of each predictor was tested at selected time points at the whole-brain level (p ≥ 0.05, FWE corrected). The table provides the peak coordinates of significant clusters, the anatomical regions contributed to significant clusters (based on the AAL Atlas), as well as beta and cluster-based t-values (df = 23). The overlap was defined as grid points contributing to both a significant effect for A_n-1_ and V_n-1_ (at any time). L: left hemisphere; R: right hemisphere. BA: Brodmann area.

| Regressor | Post-stim. time (ms) | Anatomical Labels | MNI Coord. (peak) | $\beta$ t-value |
| --- | --- | --- | --- | --- |
| | | | Brodmann Area | ($t_{sum}$) |
| $A_n$ | 80 | L: Angular, Temporal Mid. | -40, -52, 21 | 17.9 |
|  |  | L/R: Postcentral, Precuneus | BA 39 | (13862) |
|  | 190 | R: Precuneus, Lingual | 8, -51, 5 | 9.3 |
|  |  | Temporal Sup. | BA 30 | (8323) |
|  |  | L: Pre/Postcentral, Precuneus |  |  |
|  | 290 | L: Occipital Mid/Sup. | -24, -100, 5 | 6.9 |
|  |  | R: Temporal Mid/Sup., Parietal Sup. | BA 17 | (2390) |
| $A_{n-1}$ | 80 | L: Lingual, Precuneus | -24, -52, 13 | -5.8 |
|  |  | R: Lingual, Parietal Sup. | BA 17 | (-1152) |
|  | 190 | R: Frontal Mid/Inf, Precentral | 33, 21, 21 | -6.1 |
|  |  |  | BA 48 | (-1343) |
|  | 290 | R: Precuneus, Cingulum Mid/Post. | 16, -42, 41 | -4.2 |
|  |  | Parietal Inf/Sup., Postcentral | BA 23 | (-256) |
| $V_{n-1}$ | 80 | R: Fusiform, Angular, Parietal Inf. | 32, -51, -3 | 4.3 |
|  |  | Temporal Mid. Inf, Precuneus | BA 37 | (225) |
| overlap | - | R: Calcarine, Precuneus, Cingulum Mid. | 17, -67, 23 | - |
|  |  |  | BA 18 |  |

Using the third model, we directly tested whether these neural signatures of previous stimuli are significantly related to the participants’ single trial response bias (Figure 4C; Table 3). The significant influences of the neural representations of previous acoustic and visual stimuli overlapped again in parietal cortex (angular gyrus, precuneus; Figure 4C; red inset). The converging evidence from these three analyses demonstrates that the same parietal regions retain information about both previously received acoustic and visual spatial information, and that single trial variations in these neural representations directly influence the participants’ bias of subsequent sound localization.

**Table 3.** Regression Model: VAE ~ 1 + LDA_An-1_ + LDA_Vn-1_. The significance of each predictor was tested at selected time points at the whole-brain level (p≤0.05, FWE corrected). The table provides the peak coordinates of significant clusters, the anatomical regions contributing to these clusters (based on the AAL Atlas), as well as the peak beta values and cluster-based t-values (df=23). The overlap was defined as grid points contributing to both a significant effect for LDA_An-1_ and LDA_Vn-1_ (at any time). L: left hemisphere; R: right hemisphere. BA: Brodmann area. **sum of 2 spatially separate clusters, ***sum of 4 clusters

| Regressor | Post-stim. time (ms) | Anatomical Labels | MNI Coord. (peak)<br>Brodmann Area | $\beta$ t-value<br>($t_{sum}$) |
| --- | --- | --- | --- | --- |
| **$LDA_{An-1}$** | 100 | L: Occipital Mid/Sup., Temporal Inf.,<br>Parietal Mid/Sup | -32, -87, 37 | -7.4 |
|  |  | L/R: Precuneus | BA 19 | (-3571)** |
|  | 240 | L: Precentral, Frontal Mid, Precuneus,<br>Temporal Pole Sup | -32, -20, 45<br>BA 03 | -4.7<br>(-413)*** |
| **$LDA_{Vn-1}$** | 130 | R: Occipital Mid/Sup, Parietal Inf/Sup,<br>Angular | 24, -95, 29<br>BA 18 | 4.7<br>(579) |
| **Overlap ($C_{VAE}$)** | - | L/R: Precuneus | -3, -65, 51 | - |
|  |  | R: Angular | BA 07 |  |

### Neural correlates of the VE

To be able to directly compare the neural correlates of the ventriloquist aftereffect to multisensory integration within a trial (i.e. the VE effect), we repeated the same analysis strategy focusing on the MEG activity in trial n-1. As expected from the above, classification for both auditory (A_n-1_) and visual (V_n-1_) locations was significant in a network of temporal and occipital regions. We found overlapping regions in which both the acoustic and visual information significantly influenced the encoding of the response in a broad network encompassing the temporal-parietal-occipital areas during multisensory processing. To directly link the encoding of multisensory information to behavior, we again modelled the single trial VE response bias based on the representations of current acoustic and visual information (Figure 5). This revealed overlapping representations of both stimuli that directly correlated with the response bias within superior parietal regions (precuneus and superior parietal lobule), and, in a separate cluster, within inferior temporal areas (Figure 5; Table 4).

**Figure 5.**
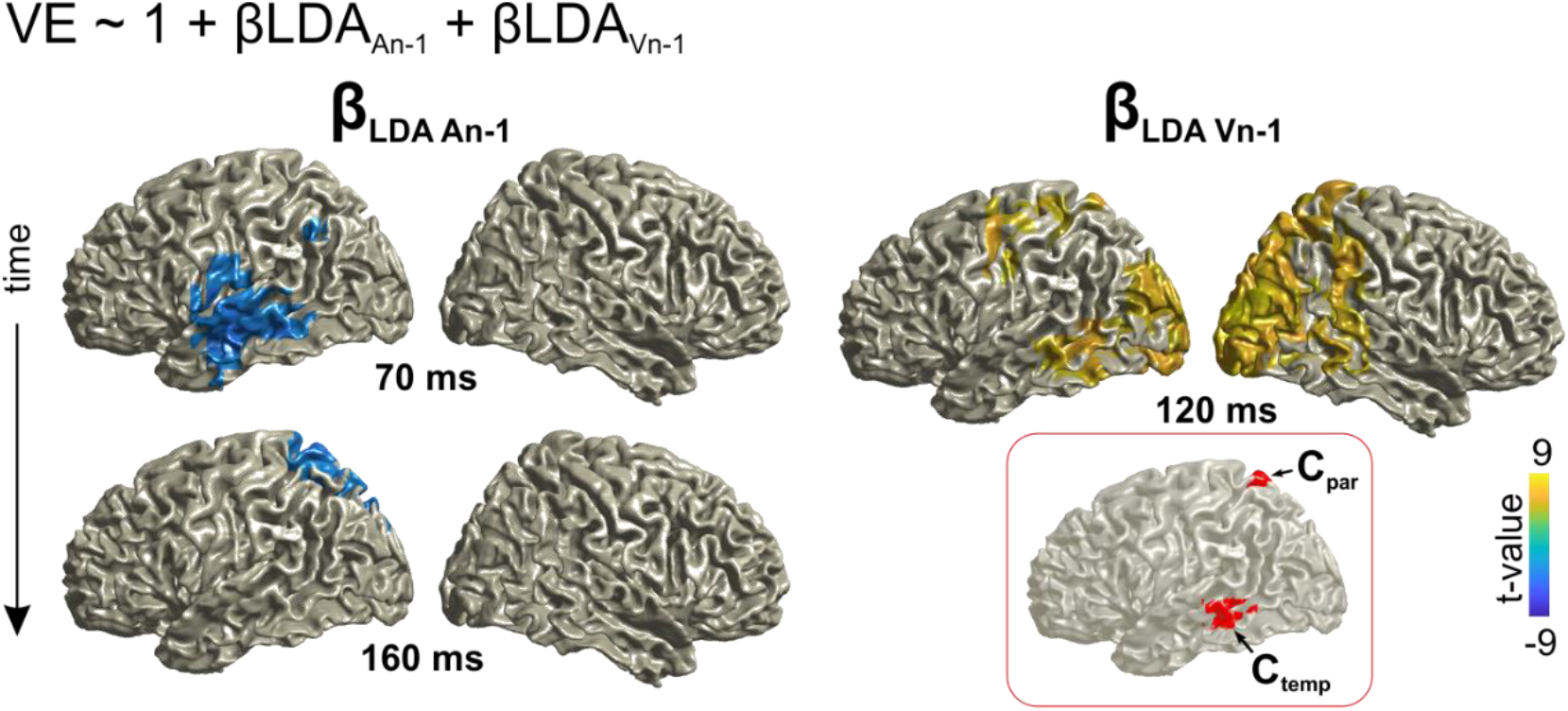
Neural correlates of the integration of auditory and visual information within a trial (VE bias). Contribution of the representations of acoustic and visual information to the single trial bias in trial n-1. (Table 4). Red inset: Grid points with overlapping significant effects for both LDAA_n-1_ and LDAV_n-1_. Surface projections were obtained from whole-brain statistical maps (at p ≤ 0.05, FWE corrected).

**Table 4.**
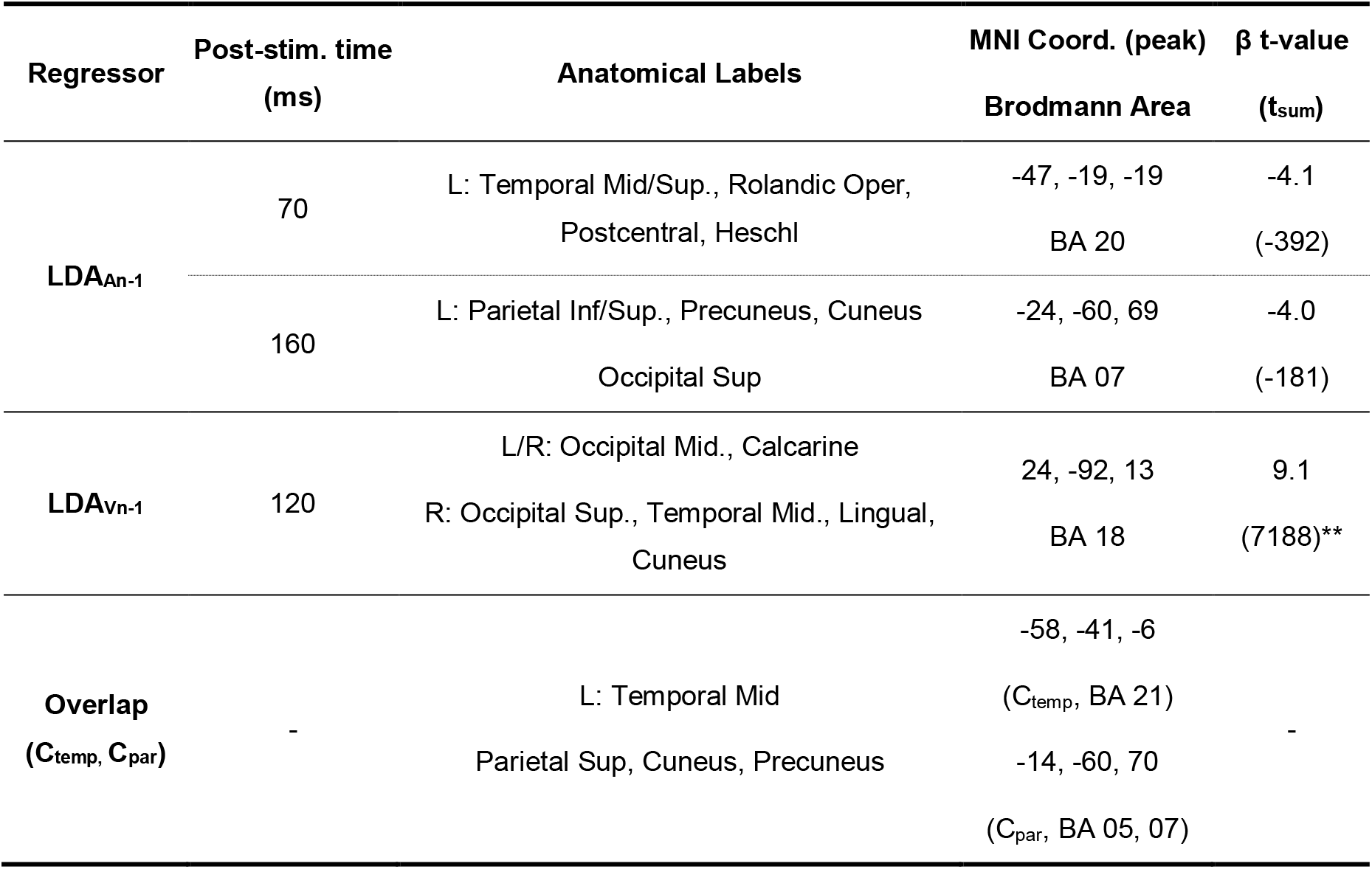
Regression Model: VE ~ 1 + LDA_An-1_ + LDA_Vn-1_. The significance of each predictor was tested at selected time points at the whole-brain level (p ≤ 0.05, FWE corrected). The table provides the peak coordinates of significant clusters, the anatomical regions contributed to significant clusters (based on the AAL Atlas), peak beta values and cluster-based t-values (df = 23). The overlap was defined as grid points contributing to both a significant effect for LDA_An-1_ and LDA_Vn-1_ (at any time). L: left hemisphere; R: right hemisphere. BA: Brodmann area. **sum of 2 spatially separate clusters,

### The same parietal regions contribute to integration within a trial and recalibration between trials

The above reveals neural representations of audio-visual information in parietal regions that either contribute to integration within a trial (VE bias) or that shape the subsequent localization of auditory information (VAE bias). Given that each effect was localized independently (as overlapping clusters in Figures 4C and Figure 5, respectively), we asked whether the same neural sources significantly contribute to both effects. To this end we subjected the above identified clusters to both neuro-behavioral models (VAE and VE; Eq. 4/5) to assess the significance of each regressor and to compare the strength of the VAE and VE effects between clusters (Table 5).

**Table 5.** Overlapping neural substrates for integration and recalibration. Both neuro-behavioral models, VE and VAE (Eq. 4/5), were tested within the clusters significantly contributing to the VAE effect (from Figure 4C, C_VAE_) and the two clusters contributing to the VE effect (from Figure 5, C_Temp_, C_par_). The table lists regression betas and group-level t-values. The expected effects (based on Figure 4C and Figure 5) are shown in normal font, the effects of interest (cross-tested) in BOLD. We directly compared the effect strengths between clusters (one-sided paired t-test, p < 0.05, FDR adjusted). Significant results are indicated by *. In particular, both C_VAE_ and C_par_ have significant VAE and VE effects (t-value), and their respective effect sizes do not differ between clusters (ns beta differences).

| model | VAE ~ 1+β*LDA<sub>An-1</sub> + β*LDA<sub>Vn-1</sub> | | | VE ~ 1+β*LDA<sub>An-1</sub> + β*LDA<sub>Vn-1</sub> | |
| --- | --- | --- | --- | --- | --- |
| cluster | $C_{VAE}$ | $C_{Temp}$ | $C_{par}$ | $C_{VAE}$ | $C_{par}$ |
| t-value | -3.29 | **-2.86** | **-2.97** | **-2.50** | -3.31 |
| ($\beta_{LDAAn-1}$) | (-0.12) | **(-0.13)<sup>ns</sup>** | **(-0.09)<sup>ns</sup>** | **(-0.30)<sup>ns</sup>** | (-0.32) |
| t-value | 3.17 | **0.91** | **2.23** | **6.61** | 6.93 |
| ($\beta_{LDAVn-1}$) | (0.12) | **(0.03)\*** | **(0.08)<sup>ns</sup>** | **(3.88)<sup>ns</sup>** | (3.35) |

This revealed that the spatially selective activity contributing to the VE effect (C_par_, from Figure 5) also significantly contributes to the VAE effect. That is, the single trial variations in the encoding of auditory and visual information in this cluster also contributed significantly (at p ≤ 0.05) to the recalibration effect. Further, the effect strength in this cluster for recalibration did not differ from that observed in the cluster directly identified as significantly contributing to the VAE bias (C_VAE_, at p < 0.05; FDR adjusted; Table 5). Vice versa, we found that the parietal sources mediating recalibration (cluster CVAE; from Figure 4C) also significantly contributed to sensory integration within trial n-1 (Table 5). These results confirm that spatially selective activity within superior parietal regions (identified by both clusters, C_par_, and C_VAE_, comprising precuneus and superior parietal regions) is significantly contributing to both sensory integration and trial-by-trial recalibration.

### Processing combined sensory evidence in multisensory clusters

The above demonstrates that medial superior parietal activity reflects both, integration and recalibration. Given that the behavioral recalibration was driven by the combined audio-visual information in the preceding trial (c.f. Figure. 2) this raises the question as to whether the neurally encoded information about the previous trial reflects previous unisensory information, or the behaviorally combined information from the previous trial. To test this, we compared the classification performance of the activity in each cluster of interest (C_VAE_, C_temp_, C_par_) for the location of the previous sound (A_n-1_) and the combined sensory information, as predicted by each participant’s behavioral weighting model (i.e., the VE bias predicted by mi_3_: β_s_*A_n-1_ + β_s_*V_n-1_, s: participant c.f. Figure. 1).

For parietal activity in trial n-1 (clusters C_VAE_ and C_par_) discriminant performance was significantly higher for the weighted multisensory than for unisensory A_n-1_ information (two-sided paired t-test, p ≤ 3*10^−6^, for both comparisons, FDR corrected; Figure 6), confirming that these regions indeed encode the integrated multisensory information. This difference was no longer significant when tested using the brain activity in trial n, possibly because the overall classification performance was lower for previous stimuli in the next trial. In contrast, temporal activity (C_temp_) was equally sensitive to unisensory and combined multisensory information in both trials.

**Figure 6.**
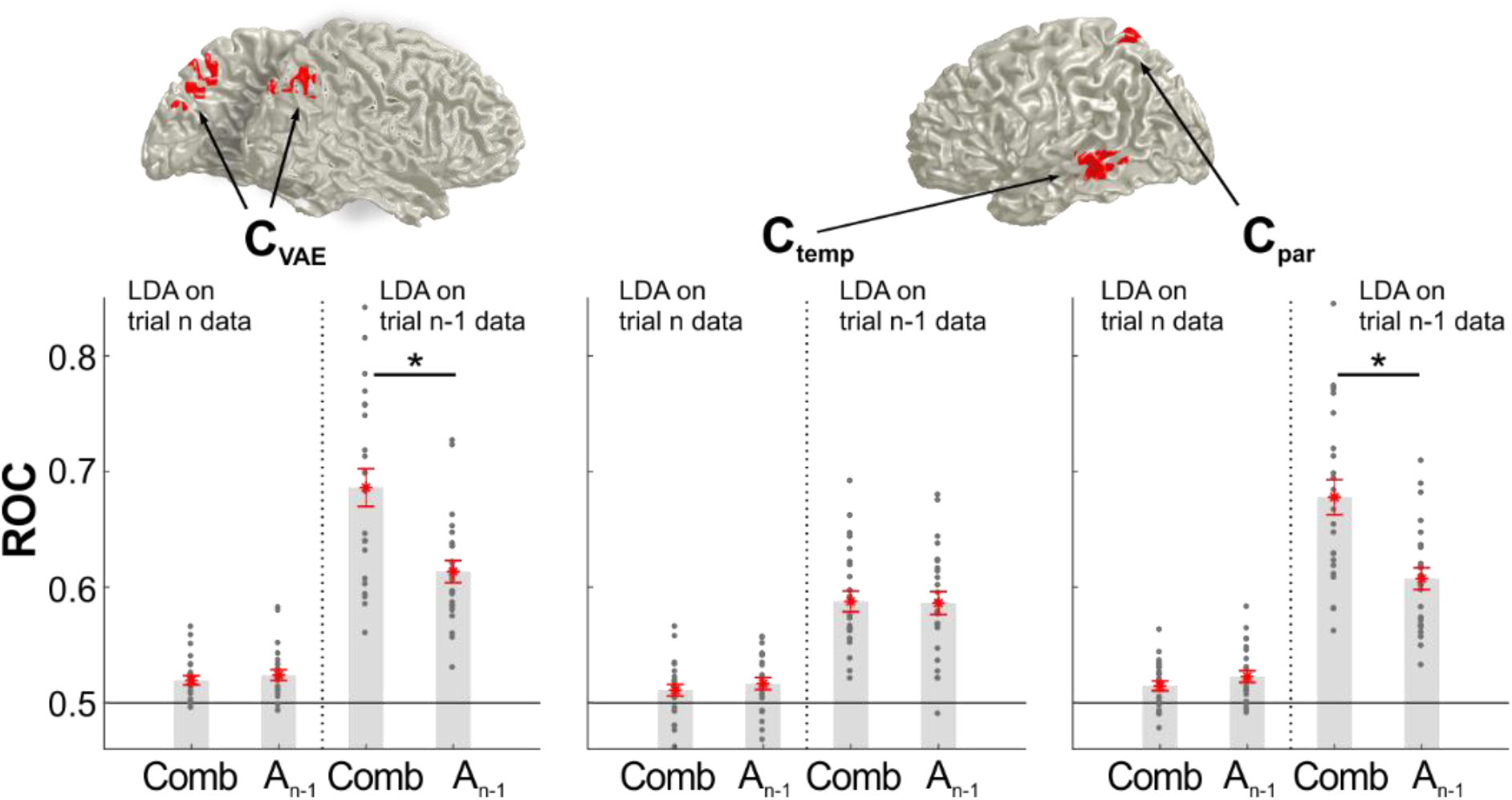
Classification performance for unisensory and combined multisensory spatial information. The bar graphs show the classification performance for each cluster of interest (C_VAE_ from Figure 4C, C_par_ and C_temp_ from Figure 5) based on the activity in trial n-1 or trial n. Classification was applied to either the sound location in trial n-1 (A_n-1_), or the combined multisensory information in trial n-1 (Comb), derived from the participant specific VE bias (derived from model mi_3_ for the behavioral data). Asterisks denote p < 0.01, two-sided paired t-test, FDR corrected for multiple comparisons at p ≤ 0.05. Gray dots are individual participant values averaged within each cluster, red stars are the mean across participants, and red lines are standard errors of mean.

## Discussion

Adaptive behavior in multisensory environments requires the combination of relevant information received at each moment in time, and the adaptation to contextual changes over time, such as discrepancies in multisensory evidence. This multi-facetted profile of flexible multisensory behavior raises the question of whether the sensory integration at a given moment and the use of previous multisensory information to recalibrate subsequent perception arise from the same or distinct neural mechanisms. We here directly compared multisensory integration and trial-by-trial recalibration in an audio-visual spatial localization paradigm. Using single trial MEG analysis we determined a network of temporal and parietal brain regions that mediate behavioral sound localization. Of these regions, superior medial parietal activity was representing current auditory and visual information, encoded the combined multisensory estimate as reflected in participant’s behavior, and retained information during the subsequent trial. Importantly, these parietal representations mediated both the multisensory integration within a trial and the subsequent recalibration of unisensory auditory perception, suggesting a common neural substrate for sensory integration and trial-by-trial recalibration of subsequent unisensory perception.

### Neural signatures of previous sensory information

Despite many behavioral studies demonstrating the robustness of multisensory trial-by-trial recalibration (6,7,9), little is known about the underlying neural substrate. Reasoning that recalibration relies on the persistence of information about previous stimuli, our study was guided by the quantification of where in the brain sensory information experienced during one trial can be recovered during the subsequent trial. This revealed persistent representations of previous acoustic information in a temporal-parietal network, suggesting that the regions known to reflect auditory spatial information within a trial also retain previous sensory evidence to mediate behavior (41–44).

Neural representations of previous visual information were also strongest in temporal and parietal regions, although classification performance was not significant at the whole brain level. Importantly, the behavioral data clearly revealed a lasting effect of visual information on behavior. Furthermore, the neuro-behavioral analysis revealed a significant contribution to behavior of neural representations of previous visual information in the superior parietal cortex. One reason for the weaker classification performance for previous visual stimuli could be a bias towards acoustic information in the participant’s task, which was to localize the sound rather than the visual stimulus in both trials. Alternatively, it could be that the visual information is largely carried by neural activity reflecting the combined audio-visual information, and hence persists directly in form of a genuine multisensory representation. This integrated multisensory representation comprises a stronger component of acoustic over visual spatial information, as reflected by the ventriloquist bias in the present paradigm. Our data indeed support this conclusion, as parietal activity within the audio-visual trial was encoding the behaviorally combined information more than the acoustic information. Previous work has shown that parietal activity combines multisensory information flexibly depending on task-relevance and crossmodal disparity (30,34) and the focus on reporting the sound location in our task may have led to the attenuation of the combined visual information in the subsequent trial. Future work is required to better understand the influence of task-relevance on uni- and multisensory representations and how these are maintained over time.

### Multiple facets of multisensory integration in parietal cortex

Several brain regions have been implied in the merging of simultaneous audio-visual spatial information (22,45,46). Our results suggest that the perceptual bias induced by vision on sound localization (i.e. the ventriloquist effect) is mediated by the posterior middle temporal gyrus and the superior parietal cortex, with parietal regions encoding the perceptually combined multisensory information. These regions are in line with previous studies, which have pinpointed superior temporal regions, the insula and parietal-occipital areas as hubs for multisensory integration (11,26,27,47). In particular, a series of studies revealed that both temporal and parietal regions combine audio-visual information in a reliability- and task-dependent manner (30,31,34,48). However, while posterior parietal regions reflect the automatic fusion of multisensory information, more anterior parietal regions reflected an adaptive multisensory representation that follows predictions of Bayesian inference models. These anterior regions (sub-divisions IPS3 and 4 in (31)) combine multisensory information when two cues seem to arise from a common origin and only partially integrate when there is a chance that the two cues arise from distinct sources, in accordance with the flexible use of discrepant multisensory information for behavior (4). Noteworthy, the peak effect for the ventriloquist aftereffect found here was located at the anterior-posterior location corresponding to the border of IPS2 and IPS3 (49), albeit more medial. Our results hence corroborate the behavioral relevance of superior-anterior parietal representations and fit with an interpretation that these regions mediate the flexible use of multisensory information, depending on task and sensory congruency, to mediate adaptive behavior

In contrast, very little is known about the brain regions implementing the trial-by-trial recalibration of unisensory perception by previous multisensory information. In fact, most studies have relied on prolonged adaptation to multisensory discrepancies. Hence these studies investigated long-term recalibration, which seems to be mechanistically distinct from the trial-by-trial recalibration investigated here (7,18,19,27,28). The study most closely resembling the present one suggested that the ventriloquist after-effect is mediated by an interaction of auditory and parietal regions (18), a network centered view also supported by work on the McGurk after-effect (14,50). Yet, no study to date has investigated the direct underpinnings of multisensory recalibration at the trial-by-trial level, or attempted to directly link the neural signature of the encoded sensory information about previous stimuli to the participant-specific perceptual bias. Our results close this gap by demonstrating the behavioral relevance of anterior medial parietal representations of previous multisensory information, which seem to reflect the flexible combination of spatial information following multisensory causal inference, and have a direct influence on participants’ perceptual bias in localizing a subsequent unisensory stimulus.

The retention of information about previous stimuli in parietal cortex directly links to animal work, which has revealed a mixed pattern of neural selectivity in parietal cortex, with individual neurons encoding both unisensory and multisensory information, reflecting the accumulation of this over time, and the transformation into perceptual choice (36). For example, parietal neurons are involved in maintaining the history of prior stimulus information, a role that is directly in line with our results (51). Our results set the stage to directly probe the correlates of multisensory recalibration at the single neuron level, for example, to address whether integration and recalibration are mediated by the very same neural populations.

In humans, the medial superior parietal cortex pinpointed here as mediator of recalibration, in particular the precuneus, has been implied in maintaining spatial and episodic memory (38–40,52,53) used for example during navigation, spatial updating or spatial search (53,54). Our results broaden the functional scope of these parietal regions in multisensory perception, by showing that these regions are also involved in the integration of multiple simultaneous cues to guide subsequent spatial behavior. This places the medial parietal cortex at the interface of momentary sensory inference and memory, and exposes multisensory recalibration as a form of implicit episodic memory, mediating the integration of past and current information into a more holistic percept.

### Integrating multisensory information across multiple time scales

Similar to other forms of memory, the history of multisensory spatial information influences perception across a range of time scales (7–9). In particular, recalibration emerges on a trial-by-trial basis, as investigated here, and after several minutes of exposure to consistent discrepancies (5,12,13,55). Behavioral studies have suggested that the mechanisms underlying the trial-by-trial and long-term effects may be distinct (9,10). Yet, it remains possible that both are mediated by the same neural mechanisms, such as the same source of spatial memory (38,52). Indeed, an EEG study on multisensory long-term recalibration has reported a neural correlates compatible with an origin in parietal cortex (19). The finding that medial parietal regions are involved in spatial and episodic memory and mediate trial by trial perceptual recalibration clearly lends itself to hypothesize that the very same regions should also contribute to long term recalibration as well.

Navigating an ever-changing world, the flexible use of past and current sensory information lies at the heart of adaptive behavior. Our results show that multiple regions are involved in the momentary integration of spatial information, and specifically expose the medial superior parietal cortex as a hub that maintains multiple sensory representations to flexibly interface the past with the environment to guide adaptive behavior.

## Methods

Twenty-six healthy right-handed adults participated in this study (13 females, age 23.7±4.3 years). Data from two participants had to be excluded as these were incomplete due to technical problems during acquisition, hence results are reported for 24 participants. All participants submitted written informed consent, and reported normal vision and hearing, and indicated no history of neurological diseases. The study was conducted in accordance with the Declaration of Helsinki and was approved by the local ethics committee (College of Science and Engineering, University of Glasgow).

### Task Design and Stimuli

The paradigm was based on an audio-visual localization task (6). Trials and conditions were designed to probe both the ventriloquist effect and the ventriloquist-aftereffect. A typical sequence of trials is depicted in Figure 1A. The participants’ task was to localize a sound during either Audio-Visual (AV) or Auditory (A), trials, or, on a subset of trials (~8% of total trials), to localize a visual stimulus (V trials). The locations of auditory and visual stimuli were each drawn semi-independently from 5 locations (−17, −8.5, 0, 8.5, 17 °of visual angle from the midline), to yield 9 different audio-visual discrepancies (−34, −25.5, −17, −8.5, 0, 8.5, 17, 25.5, 34 °). Importantly, AV and A trials were always presented sequentially to probe the influence of audio-visual integration on subsequent unisensory perception. In the following we refer to the AV trials also as trial n-1, and the subsequent A trial as trial n, to explicitly denote their temporal sequence (Figure 1A).

Acoustic stimuli were spatially dispersed white noise bursts, created by applying head related transfer functions (HRTF) (PKU & IOA HRTF database, Qu et al., 2009) to white noise (duration = 50 ms) defined at specific azimuths, elevations and distances. Here we used a distance of 50 cm and 0 elevation. Sounds were sampled at 48 kHz, delivered binaurally by an Etymotic ER-30 tube-phone at ~84.3 dB (root-mean-square value, measured with a Brüel & Kjær Type 2205 sound-level meter, A-weighted). An inverse filtering procedure was applied to compensate for the acoustic distortion introduced by plastic tubes required for the use in the MEG shield-room (57). The visual stimulus was a white Gaussian disk of 50 ms duration covering 1.5° of visual angle (at full-width half-maximum). This was back-projected onto a semi-transparent screen located 50 cm in front of the participant, via a DLP projector (Panasonic D7700). Stimulus presentation was controlled with Psychophysics toolbox (Brainard, 1997) for MATLAB (The MathWorks, Inc., Natick, MA), with ensured temporal synchronization of auditory and visual stimuli.

In total we repeated each discrepancy (within or between trials) 40 times, resulting in a total of 360 AV-A trial pairs. In addition, 70 visual trials were interleaved to maintain attention (V trials always came after A trials, thus not interrupting the AV-A pairs), resulting in a total of 790 trials (2 × 360 + 70) for each participant. Trials were pseudo-randomized, and divided into 10 blocks of ~ 8 mins each. Each trial started with a fixation period (uniform range 800 – 1200 ms), followed by the stimulus (50 ms). A response cue was presented after a random interval (uniform range 600 – 800 ms) and participants responded by moving a trackball mouse (Current Designs Inc., Philadelphia, PA 19104 USA) with their right hand by moving the cursor to the location of the perceived stimulus and clicking the button. A letter ‘S’ was displayed on the cursor for ‘sound’, and ‘V’ for the visual trials. There was no constraint on response times. Inter-trial intervals varied randomly (uniform 1100-1500 ms). The experiment lasted about 3.5 hours including preparation and breaks.

### Analysis of behavioral data

#### Ventriloquist effect (VE)

For AV trials, we defined the VE as the difference between the reported location (R_n-1_) and the location at which the sound (A_n-1_) was actually presented (R_n-1_ – A_n-1_). To determine whether the response bias captured by the VE was systematically related to any of the sensory stimuli, we compared the power of different linear mixed models for predicting this responses bias as computational accounts for the VE. These models relied either on only the auditory, only the visual stimulus location, or their combination: mi_1_: VE ~ 1 + β·A_n-1_ + subj, mi_2_: VE ~ 1 + β·V_n-1_ + subj, mi_3_: VAE ~ 1 + β·A_n-1_ + β·V_n-1_ + subj, where A_n-1_ and V_n-1_ were the main effects, and the subject ID was included as random effect. Models were fit using maximum-likelihood procedures and we calculated the relative Bayesian information criterion (BIC) (BIC -mean(BIC_m_), and relative Akaike information criterion (AIC) (AIC – mean(AIC_m_)), and the protected exceedance probability (58) for formal model comparison.

#### Ventriloquist-aftereffect (VAE)

The VAE was defined as the difference between the reported location (R_n_) minus the mean reported location for all trials of the same stimulus position. This was then expressed as a function of the audio-visual discrepancy in the previous trial (i.e., V_n-1_-A_n-1_). Again we used linear mixed-effects models to compare different accounts of how the response bias depends on the stimuli: mr_1_: VAE ~ 1 + β·A_n-1_ + subj, mr_2_: VAE ~ 1 + β·V_n-1_ + subj, mr_3_: VAE ~ 1 + β·A_n-1_ + β·V_n-1_ + subj. We also quantified the response bias as a function of the all stimulus locations (i.e. A_n_, V_n-1_, A_n-1_), in order to determine the influence of each individual stimulus on behavior (Figure 1C).

### Magnetoencephalography (MEG) acquisition

Participants were seated in a magnetically shielded room, 50 cm in front of a screen (30.5 cm × 40.5, 1024 × 768 resolution). The MEG data was recorded with a 248-magnetometer, whole-head MEG system (MAGNES 3600 WH, 4-D Neuroimaging, San Diego, CA) at a sampling rate of 1017.25 Hz. Head positions were measured at the beginning and end of each block, using 5 coils marking fiducial landmarks on the head of the participants, to monitor head movements. Coil positions were co-digitized with the head shape (FASTRAK, Polhemus Inc., Colchester, VT).

### Preprocessing of MEG data

MEG data were preprocessed with MATLAB (The MathWorks, Inc., Natick, MA) using the Fieldtrip toolbox (version 20171001, Oostenveld, Fries, Maris, & Schoffelen, 2011). Each block was preprocessed individually (ft_preprocessing). Epochs from −0.6 ~ 0.6 (0 = stimulus onset) were extracted from the continuous data, and denoised using the MEG reference (ft_denoise_pca). Resulting data was filtered between 1~48 Hz (4-order Butterworth filter, forward and reverse), and down-sampled to 100 Hz. Known faulty channels (N = 3) were removed. Then, variance, maximum, minimum, and range of data across trials were calculated for each channel, and channels with extreme data were excluded. Outliers were defined based on interquartiles (IQR) (60); Q1 – 4.5×IQR or above Q3 + 4.5×IQR. Weight 4.5 instead of the standard 1.5 was used since 1.5 was eliminating too many channels. Overall about 6% of all channels were excluded (15.1±5.8 channels per participant; mean±SD). Heart and eye-movement artifacts were removed using independent component analysis (ICA) with Fieldtrip (ft_componentanalysis, ft_rejectcomponent), which was calculated based on 30 principal components. Trials with SQUID jumps were detected and removed (ft_artifact_jump) with a cutoff z-value of 20. Finally, the data was manually inspected using the interquartile method (across channels weight 2.5) to exclude outlier trials. On average about 2% of trials had to be discarded (18.3±7.7 trials per participant; mean±SD, including very few trials on which the mouse button did not react properly).

### MEG Source Reconstruction

Source reconstruction was performed using Fieldtrip (59), SPM8 (Wellcome Trust, London, United Kingdom), and the Freesurfer toolbox (61). For each participant whole-brain T1-weighted structural magnetic resonance images (MRIs, 192 sagittal slices, 256 × 256 matrix size, 1 mm^3^ voxel size) were acquired using a Siemens 3T Trio scanner (32-channel head coil). These were co-registered to the MEG coordinate system using a semi-automatic procedure. Individual MRIs were segmented and linearly normalized to a template brain (MNI space). Next, a volume conduction model was constructed using a single-shell model based on an 8 mm isotropic grid. We projected the sensor-level waveforms into source space using a linear constraint minimum variance (LCMV) beamformer with a regularization parameter of 7%. Then the data was collapsed onto the strongest dipole orientation based on singular value decomposition. Source reconstruction was performed on each block separately, and then concatenated for further analyses.

### Discriminant analysis

To extract neural signatures of the encoding of different variables of interest we applied a cross-validated regularized linear discriminant analysis (LDA) (62,63) to the single trial MEG source data. LDA was applied to the data aligned to stimulus onset in 60 ms sliding windows, with 10 ms time-steps, using a spatial searchlight around each voxel consisting of the 27 neighboring voxels. For each source point *ν*, the LDA identifies a projection of the multidimensional source data, *x_ν_*(*t*), that maximally discriminates between the two conditions of interest, defined by a weight vector, *w_ν_*, which describes a one dimensional combination of the source data, *Y_ν_*(*t*):

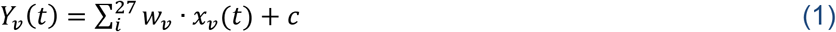

with *i* summing over grid points within a spatial searchlight, and *c* being a constant. Classification performance was quantified using the area under the receiver operator characteristic (ROC; Az value) based on 6-fold cross validation. We identified clusters with significant classification performance at the group level by applying a cluster-based permutation procedure (see below). Having established a set of discriminant weights, *w_ν_*, one can derive single trial predictions of the neurally encoded information, *Y_ν_*(*t*), using equation 1. Importantly, using cross-validation one can determine the classification weights on one set of trials, and then predict the discrimination performance for a separate trials. The value, *Y_ν_*(*t*), of such an LDA projection can serve as a proxy to the neurally encoded signal trial information about a specific stimulus variable, and can be related e.g. to behavioral performance (64,65) (64–67).

### Neural en- and decoding analysis

We computed separate linear discriminants for the location of the auditory and visual stimuli in trial n-1 (A_n-1_, V_n-1_), the auditory stimulus in trial n (A_n_), and the responses in either trial (R_n-1_, R_n_). Each location was considered as a binary variable, that is, e.g. whether the stimulus was to the left (−17, −8.5 °) or right (17, 8.5 °) of the fixation point. To elucidate neural mechanisms of the VE and VAE, we investigated different models capturing potential influences of each stimulus on the neural representation of sensory information, or on the neural representation of the upcoming participant’s response. In addition, we investigated models directly capturing the neuro-behavioral relation between the encoded sensory information and the response bias.

First, we determined when and where neural signatures of the encoding of single trial information, as reflected by their LDA discriminant values (c.f. Eq. 1), were influenced by the sensory stimuli in the current and/or previous trial. For each searchlight and time point we determined the following models:

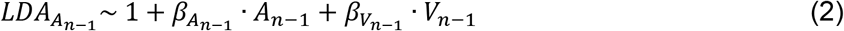

which captures sensory integration within the AV trials, and,

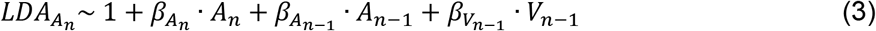

which captures how the encoding of the current sound in the A trial is influenced by previous audio-visual stimuli. Second, and analogously, we investigated models for the encoding of the participant’s response (i.e. LDA_Rn_ and LDA_Rn-1_).

Third, we determined the contribution of the single trial representations of acoustic and visual information to the single trial behavioral response bias with the following models:

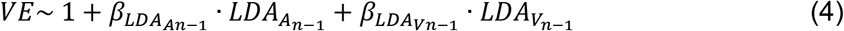

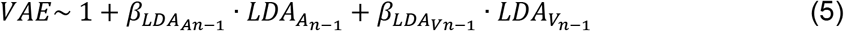

To avoid overfitting we computed these models based on 6-fold cross-validation, using distinct sets of trials to determine the weights of the LDA and to compute the regression models. We then computed group-level t-values for the coefficients for each repressor at each grid and time point, and assessed their significance using cluster-based permutation statistics (below). Note that for AV trials, we excluded the AV pairs with the most extreme discrepancies (±34 °), as these were inducing strong correlations between the regressors.

### Statistical Analysis

To test the significance of the behavioral biases we used two-sided Wilcoxon signed rank tests, correcting for multiple tests using the Holm procedure with a family-wise error rate of p= 0.05.

Group-level inference on the 3-dimensional MEG source data was obtained using randomization procedures and cluster-based statistical enhancement controlling for multiple comparisons in source space (68,69). First, we shuffled the sign of the true single-subject effects (the signs of the chance-level corrected Az values; the signs of single-subject regression beta’s) and obtained distributions of group-level effects (means or t-values) based on 2000 randomizations. We then applied spatial clustering based on a minimal cluster size of 6 and using the sum as cluster-statistics. For testing the LDA performance, we thresholded effects based on the 99th percentile of the full-brain distribution of randomized Az values. For testing the betas in regression models, we used parametric thresholds corresponding to a two-sided p=0.01 (tcrit = 2.81, d.f. = 23; except for the analysis for the visual location in trial n-1, not shown). The threshold for determining significant clusters for classification performance was p≤0.01 (two-sided), that for significant neuro-behavioral effects (Eq. 2–5) p ≤ 0.05 (two-sided). To simplify the statistical problem, we tested for significant spatial clusters at selected time points only. These time points were defined based on local peaks of the time courses of respective LDA ROC performance (for models in Eq. 2/3), and the peaks for the beta time-course for the behavioral models (Eq. 4/5). Furthermore, where possible, to test for the significance of individual regressors we applied a spatial a priori mask derived from the significance of the respective LDA ROC values to further ensure that neuro-behavioral effects originate from sources with significant encoding effects (70). Note that this was not possible for LDA_Vn-1_ for which we used the full brain to test for significant model effects.

## Acknowledgements

This work was supported by the European Research Council (to C.K. ERC-2014-CoG; grant No 646657). We would like to thank Yinan Cao & Bruno Giordano for helping with the acoustic stimuli, Bruno Giordano for helpful discussions and Gavin Paterson for support with hardware and data acquisition training along with Frances Crabbe.

